# Antibiotic treatment and microbiome depletion slow cnidarian regeneration

**DOI:** 10.1101/2024.11.24.625057

**Authors:** Jeric Da-Anoy, Kyle Toyama, Oliwia Jasnos, Audrey Wong, Thomas D. Gilmore, Sarah W. Davies

**Affiliations:** Department of Biology, Boston University, Boston, Massachusetts, USA 02215

**Keywords:** microbiome, Aiptasia, Nematostella, antibiotic, regeneration, gnotobiotic

## Abstract

Cnidarian microbiomes play an essential role in physiology and development, but how these communities influence tissue regeneration is poorly understood. Here, we examined the effects of antibiotic exposure on regeneration and microbial communities in two cnidarian models, the sea anemones *Nematostella vectensis* (non-symbiotic, hereafter, *Nematostella*) and *Exaiptasia pallida* (symbiotic, hereafter, *Aiptasia*). Bisected animals were incubated in either sterile or antibiotic-containing artificial seawater for seven days and regeneration was monitored daily. After seven days, tentacle number and length were measured, and microbial communities were profiled using metabarcoding of the V4 region of the 16S rRNA. Microbiome disruption was observed under antibiotic treatment in both species, resulting in decreased microbial load and shifts in relative abundances of some microbial taxa. However, *Nematostella* exhibited a greater reduction in microbial diversity and community shifts under antibiotic exposure, whereas *Aiptasia* showed only moderate changes in diversity. In both species, microbiome disruption was associated with slower regeneration rates and reduced tentacle number and length, suggesting a functional role for the microbiome in cnidarian regeneration. Our findings suggest that host-microbiome interactions in both symbiotic and aposymbiotic anemones are important for the maintenance of regenerative processes. These findings provide insight into how cnidarians and their microbiomes respond to environmental stressors, with implications for predicting cnidarian resilience in the context of emerging threats to the marine environment.

## Introduction

Microbiomes play essential roles in the health and resilience of coral reef ecosystems (reviewed in (Rädecker et al. 2015)). These microbial communities form complex symbioses with their cnidarian hosts that can influence health, development, and survival outcomes (MacVittie et al. 2024). For example, some bacteria associated with a cnidarian host are involved in nitrogen cycling, which is crucial for the nutritional balance within coral reefs (Ceh et al. 2013). Additionally, microbiomes can modulate immune systems, protect against pathogens, and influence metabolic processes of their cnidarian hosts (Krediet, Ritchie, et al. 2013).

Several studies have demonstrated how symbiosis can influence cnidarian immunity. Studies in Hydra have shown that symbiotic algae can modify host immune responses through the differential expression of antimicrobial peptides (Franzenburg et al. 2013). Furthermore, comparative analyses of starvation responses in non-symbiotic sea anemones (e.g., *Nematostella vectensis*) and symbiotic cnidarians (such as the anemone *Exaiptasia pallida* and some reef-building corals) have revealed distinct defense responses (Valadez-Ingersoll et al. 2024). These defense and immune pathways are modulated through complex signaling that optimizes mutualistic benefits, thereby enhancing host resilience and imposing adaptive demands on the immune system (Detournay et al. 2012; Mydlarz et al. 2016). Taken together, such studies indicate that symbiotic relationships may drive evolutionary innovations in immune systems (Davy et al. 2012).

While interactions between cnidarians and their endosymbiotic algae are well-studied, the roles of cnidarian holobiont members—which include host animals, Symbiodiniaceae algae, and associated bacteria—are increasingly being recognized for supporting stress responses and host development. For instance, microbiomes in the mucus protect cnidarian hosts from opportunistic pathogens through competitive niche occupation (Krediet, Ritchie, et al. 2013), disruption of pathogen establishment (Ceh et al. 2013), inhibition of quorum sensing (Kalia et al. 2018), and the production of antimicrobial compounds (Nissimov et al. 2009). The microbiome is also essential for cnidarian development (Thompson et al. 2014), where disruptions in bacterial composition lead to altered development (Klein et al. 2021). Exposure of juvenile *Nematostella* anemones to antibiotics leads to shifts in microbial communities, and *Nematostella* larvae exposed to antibiotics show a 50% delay in settlement time (Krueger et al. 2024). Moreover, adult *Nematostella* exposed to acute and chronic antibiotic treatments showed differential expression of development-associated genes (Krueger et al. 2024). In Hydra, the elimination of a microbiome leads to increased formation of ectopic tentacles when animals are exposed to alsterpaullone, a chemical that activates the Wnt-signaling pathway (Taubenheim et al. 2020). Additionally, interactions between pathogenic *Turneriella* species and beneficial *Pseudomonas* species in Hydra have been shown to disrupt tissue homeostasis, resulting in tumors and additional tentacles (Rathje et al. 2020). Microbiome disruption has also been linked to reduced *Aiptasia* fitness, including decreased asexual reproduction, reduced biomass, and loss of algal symbionts (MacVittie et al. 2024).

Not all cnidarian-associated bacteria are beneficial. Certain members of the Vibrio family can infect coral tissues and cause disease (Kushmaro et al. 1996; Ben-Haim et al. 2003; Rosenberg and Falkovitz 2004). Recently, antibiotics have been used to combat pathogens causing coral diseases (Hartman et al. 2022; Studivan et al. 2023). While these treatments can effectively halt disease progression and protect affected colonies, they also disrupt the natural microbial balance within hosts. Antibiotics can lead to altered gene expression, decreased photosynthetic capacity, increased reactive oxygen species (ROS) production, and subsequent cellular stress (Motone et al. 2020). Furthermore, antibiotic treatment can result in microbiome shifts, allowing opportunistic bacteria and pathogens to invade due to the elimination of dominant, protective bacteria (Sweet et al. 2011; MacVittie et al. 2024). These shifts can increase necrosis and mortality rates, as the loss of protective microbes renders the host more susceptible to harmful pathogens (Bent et al. 2021).

Previous work has shown that after antibiotic treatment, coral microbiomes can recover to a state similar to pre-treatment conditions when given sufficient time (Sweet et al. 2011), likely due to incomplete elimination of bacteria under antibiotic treatment. However, long-term effects of antibiotic use in marine ecosystems are not fully understood, and there is a need to explore how antibiotic treatment influences fitness-related traits. Here, we determined the effects of antibiotic treatment on cnidarian microbiomes and how this treatment impacted two key fitness traits: regeneration and growth. We conducted experiments on two sea anemones, *Nematostella* (non-symbiotic) and *Aiptasia* (facultatively symbiotic), both of which are model organisms commonly used in microbiome and/or symbiosis research due to ease of culture and ability to reproduce sexually and asexually (Baumgarten et al. 2015). Bisected anemones were exposed to an antibiotic cocktail and monitored regeneration rates (ability of animals to regrow tentacles after horizontal bisection of the body) and tentacle growth (number and length of new tentacles). Antibiotic treatment altered microbiome profiles, and also caused slower regeneration rates and reduced tentacle growth. These results shed light on host-microbe interactions and their broader implications for cnidarian health.

## Methods

### Animal preparation

Individual animals were acclimated separately in sterile 60 x 15 mm plates containing sterile artificial seawater (ASW) at 24°C in an incubator for one week. *Nematostella vectensis* (20– 25mm oral disk) strains from Maine (ME) and North Carolina (NC) were reared in ASW at 11 ppt, while *Aiptasia* strains CC7 (Sunagawa et al., 2009) and H2 (Xiang et al., 2013) (20–25mm oral disk) were reared in ASW at 33 ppt. A total of 150 *Nematostella* (75 individuals/strain) and 150 *Aiptasia* (75 individuals/strain) were used. Animals were fed 48 hour-old brine shrimp (*Artemia salina*) four days prior to experimental onset. The incubator was illuminated under low photosynthetic photon flux density (∼50 µmol m^−2^ s^−1^) on a 12:12 light-dark cycle.

### Antibiotic treatment preparation

Ampicillin (Amp, Acros Organics Cat. No.: 61177), streptomycin (Strep, Alfa Aesar, Cat. No.: J61299), and neomycin (Neo, Alfa Aesar, Cat. No.: J61499) were dissolved in sterile deionized water to a working concentration of 20 mg/ml. Rifampicin (Rif, Tokyo Chemical Industry, Cat. No.: 236-312-0) was dissolved in dimethyl sulfoxide (DMSO) to 20 mg/ml. As a vehicle control, DMSO was added to seawater at equal concentration to antibiotic treatment (0.1 and 0.4%, v/v). The antibiotic cocktail was a mixture of four antibiotics at concentrations of 200 µg/ml for each Amp+Strep+Neo and 50 µg/ml for Rif. All antibiotics were dissolved in ASW with identical salinity used to maintain animals.

### Antibiotic experiments

Regeneration experiments were performed on adult animals (20–25mm oral disk). Using a sterile no. 10 disposable scalpel, animals were bisected at the base of the pharynx, and only aboral physa were used (Fig. 1a). Animals were placed into separate sterile petri dishes (60 mm × 15 mm) using disposable pipettes, and maintained at 24°C in an incubator with light levels of ∼50 µmol m^−2^ s^−1^ on a 12:12 light-dark cycle. Animals were subjected to one of two treatments: 1) control, sterile ASW; 2) antibiotic cocktail. All ASW had the same salinity used to maintain the animals. Culture media was replaced daily, and animals were not fed during regeneration experiments. After 7 days, all anemones regardless of regeneration state were flash-frozen in liquid nitrogen for DNA extraction and microbiome profiling.

**Fig. 1.**
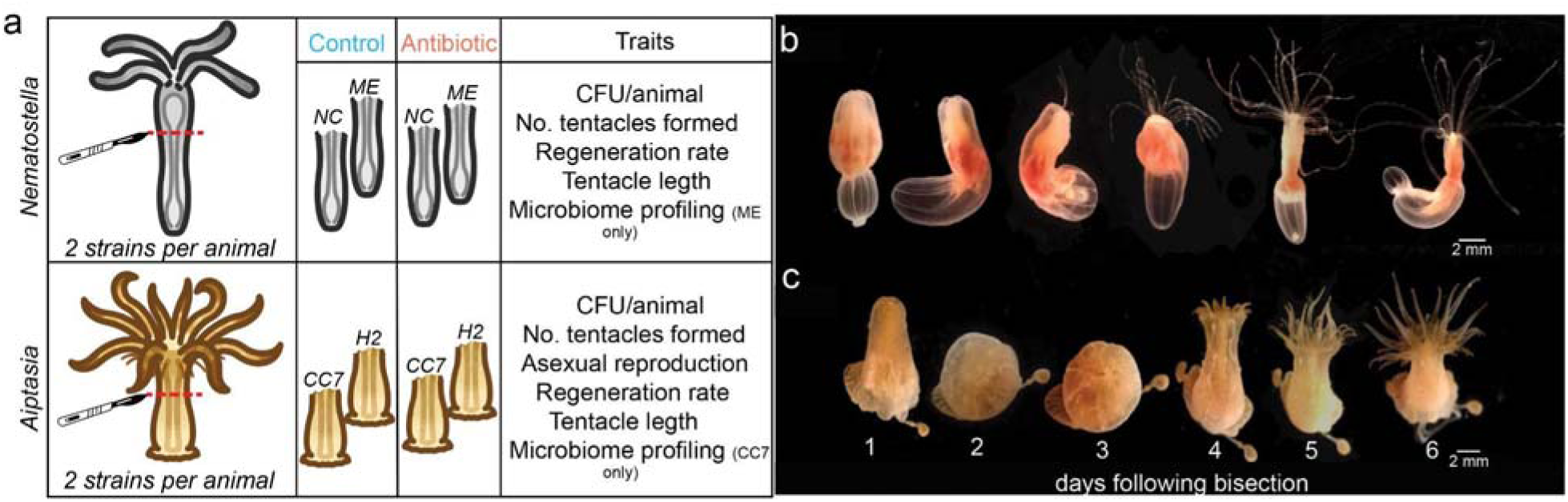
(a) Schematic of the experimental design. Two strains of *Nematostella* and *Aiptasia* were bisected and subjected to control (blue) or antibiotic (red) treatment. Representative images showing regeneration stages in *Nematostella* (b) and *Aiptasia* (c) in controls. Numbers correspond to days following bisection. Tentacle reemergence was observed after 3 and 4 days in *Nematostella* and *Aiptasia*, respectively.

### Bacterial load quantification

To quantify bacterial load, colony forming unit (CFU) assays were performed on whole animal lysates. Individual anemones (8 replicates/treatment/strain/animal) were rinsed with sterile ASW, homogenized in sterile phosphate-buffered saline (PBS) and homogenate was serially diluted in sterile PBS up to 10^8, and 10 µl of each dilution was spot plated onto Marine Agar plates. Plates were incubated at 28°C for 48–72 h, and then colonies were counted. Negative controls (sterile ASW, 1X PBS) were included to confirm absence of contamination during sample preparation.

### Regeneration rate, asexual reproduction, and tentacle phenotypes

Daily photographs of animals were taken using an Olympus Tough TG-5, with a ruler included as a size standard. Animals were scored as: 1 = regenerated or 0 = not regenerated where regeneration was identified by reemergence of the oral disc and primary tentacle regrowth (Fig. 1a). On the final day, mean number of tentacles and lengths of the five longest tentacles per animal were quantified in ImageJ. To assess the effects of antibiotics on asexual reproduction, we counted new individuals produced by longitudinal fission (splitting along body) in *Nematostella* and through pedal laceration (tissue fragments detach from pedal disc and develop into new individuals) in *Aiptasia* at the end of the experiment. Notably, asexual reproduction was not observed in *Nematostella* in either treatment.

### Statistical analyses

To estimate differences in regeneration rates between antibiotic and control treatments, *survival* v.3.5-5 and *survminer* v.0.4.9 (Therneau and Lumley 2015) packages in R version 4.4.0 were used. We compared regeneration curves using a log-rank test and assessed significant differences between treatments. Differences in CFU, number of clones produced, tentacle number and length among treatments were analyzed using an ANOVA (*aov*) with fixed effects of treatment and strain, after checking for normality with the Shapiro-Wilk Test (*shapiro.test*). Post-hoc pairwise comparisons were conducted using Tukey’s HSD tests. Results are reported in Table 1. Raw data and code for all analyses can be found on GitHub (https://github.com/jpdaanoy2024/Proj_Regeneration). Sequence reads can be accessed in the NCBI short read archive (SRA) database under the BioProject accession number PRJNA1165229.

**Table 1.**
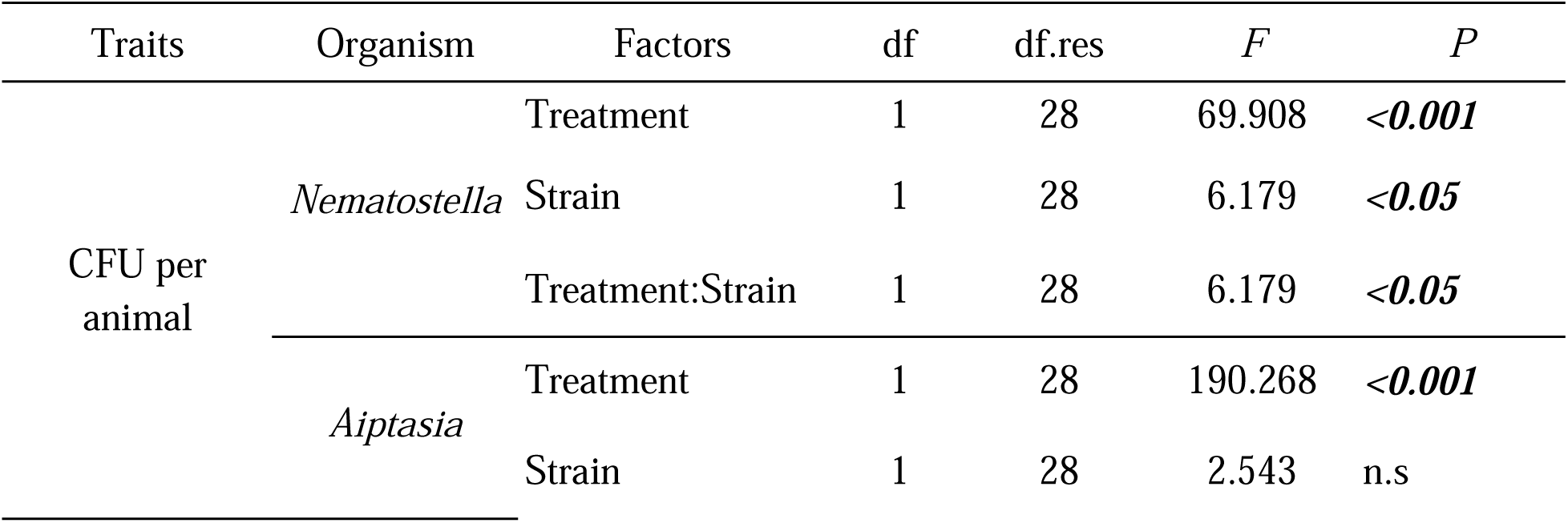
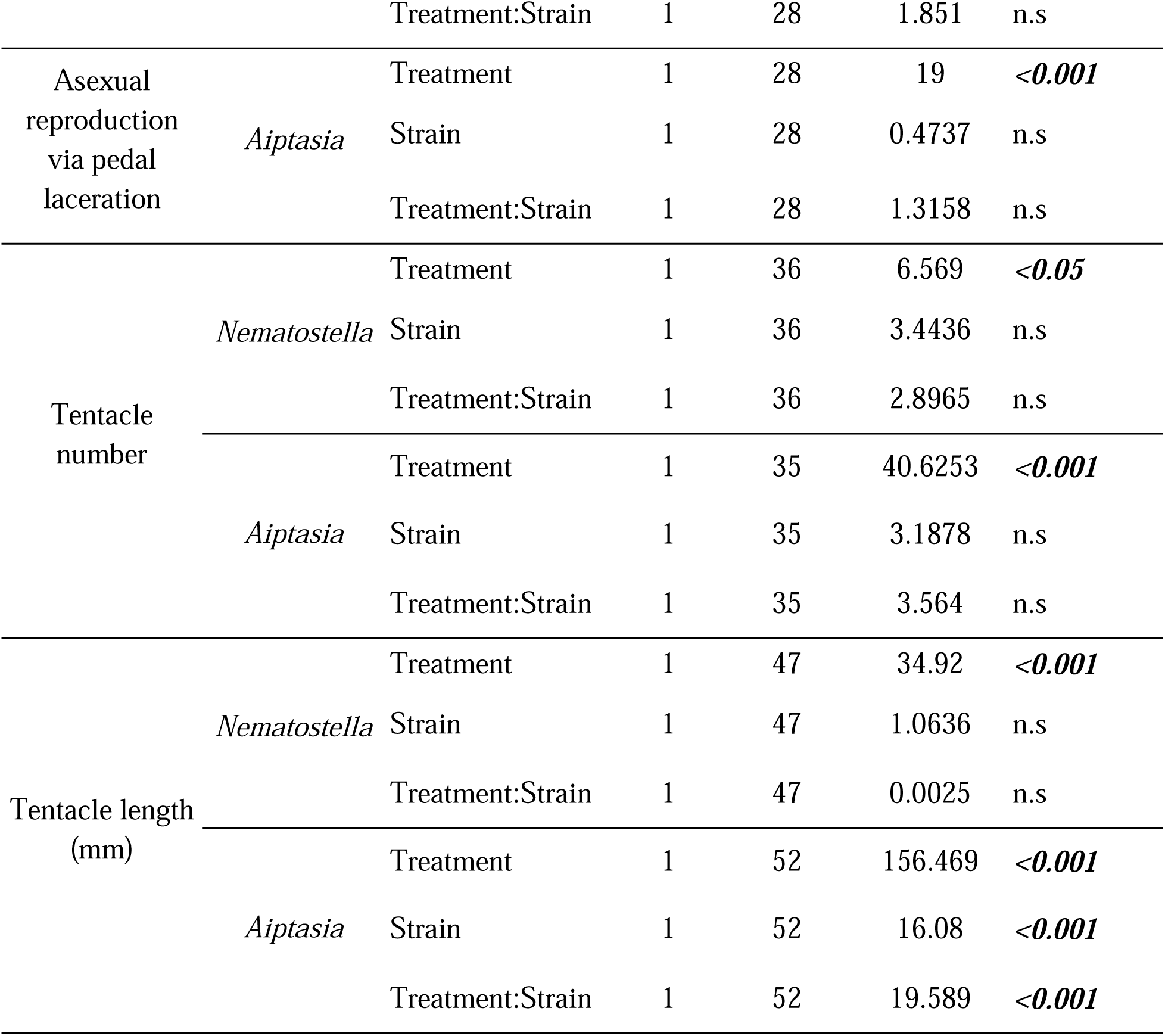
Summary of statistical results of microbial load and tentacle phenotypes (number and length) in *Nematostella* and *Aiptasia* across different treatments and strains.

### Microbiome profiling

Genomic DNA (gDNA) was isolated from each anemone (n=4-5/treatment/species) by homogenizing samples using 1 mm glass beads (BioSpec) followed by extraction with phenol:chloroform:isoamyl alcohol (25:24:1). Only one strain from each species was used: ME for *Nematostella* and CC7 for *Aiptasia*. The V4 region of the 16S rRNA was amplified from gDNA using the Hyb515F (Parada et al. 2016) and Hyb806R (Apprill et al. 2015) primers. The PCR mix contained 0.1 µl of gDNA template, 0.025 U of ExTaq, 1x ExTaq buffer (Takara), 1 µM of primers, 0.2 mM of dNTPs, and molecular biology grade water. PCR conditions were as follows: denaturation at 95°C for 5 min, followed by 35 cycles of 95°C for 1 min, 62°C for 2 min, 72°C for 2 min, and a final elongation step at 72°C for 10 min. Samples were then purified using the GeneJet PCR Purification Kit (Thermo Fisher) and subjected to a second 6-cycle PCR for dual-indexing, using identical PCR conditions. Samples were pooled and sequenced at Tufts University Core Facility on an Illumina MiSeq (250 bp paired-end).

Demultiplexed reads were pre-processed using *bbmap* (Bushnell 2014) to retain sequences containing 16S primers only. Primers were removed with *cutadapt* (Martin 2011). Quality filtering of amplicon sequence variants (ASVs) and taxonomic assignment were performed using *DADA2* (Callahan et al. 2016) with the Silva v.138.1 database (Quast et al. 2013) and the National Center for Biotechnology Information nucleotide database via blast+ (Camacho et al. 2009). Sequence reads from chloroplast, mitochondrial, non-bacterial, and negative control contaminants were removed using *decontam* (Davis et al. 2018). Rarefied ASVs were trimmed with *MCMC.OTU* (Green et al. 2014) and further processed to eliminate ASVs with low counts (<0.01%) using *vegan* (Oksanen et al. 2013).

Alpha diversity indices (Shannon index, Simpson’s index, ASV richness, and evenness) were calculated on rarefied counts using the *estimate richnes*s function in *Phyloseq* (McMurdie and Holmes 2013). Community distance matrices based on Bray-Curtis dissimilarity were assessed using principal coordinates analysis (PCoA) with *Phyloseq* and permutational multivariate analysis of variance (PERMANOVA, *vegan* package) to evaluate changes in microbiome structure and composition across treatments. All read tracking statistics are provided in Table S2.

## Results

### Reduced microbial load in anemones after exposure to antibiotics

To determine the effect of antibiotic treatment on bacterial load, CFU assays quantified culturable bacteria on marine agar. Results showed that antibiotic treatment resulted in a 98.4% reduction in CFU per individual in *Nematostella* (Fig. 2a, F(1,28)=69.91, *p<0.001*) relative to the untreated control. In addition, strain ME *Nematostella* had higher bacterial loads under control conditions compared to the NC strain and although both strains exhibited strong bacterial load reductions, a treatment *x* strain interaction (*p<0.05*) was observed, indicating that ME anemones exhibited a greater microbial load reduction following antibiotic treatment. Similarly, antibiotic treatment led to a 96.9% reduction in CFU in *Aiptasia* (Fig. 2a, F(1,28) =190.27, *p<0.001*) relative to controls, with CC7 and H2 showing similar levels of microbial depletion following antibiotic treatment.

**Fig. 2.**
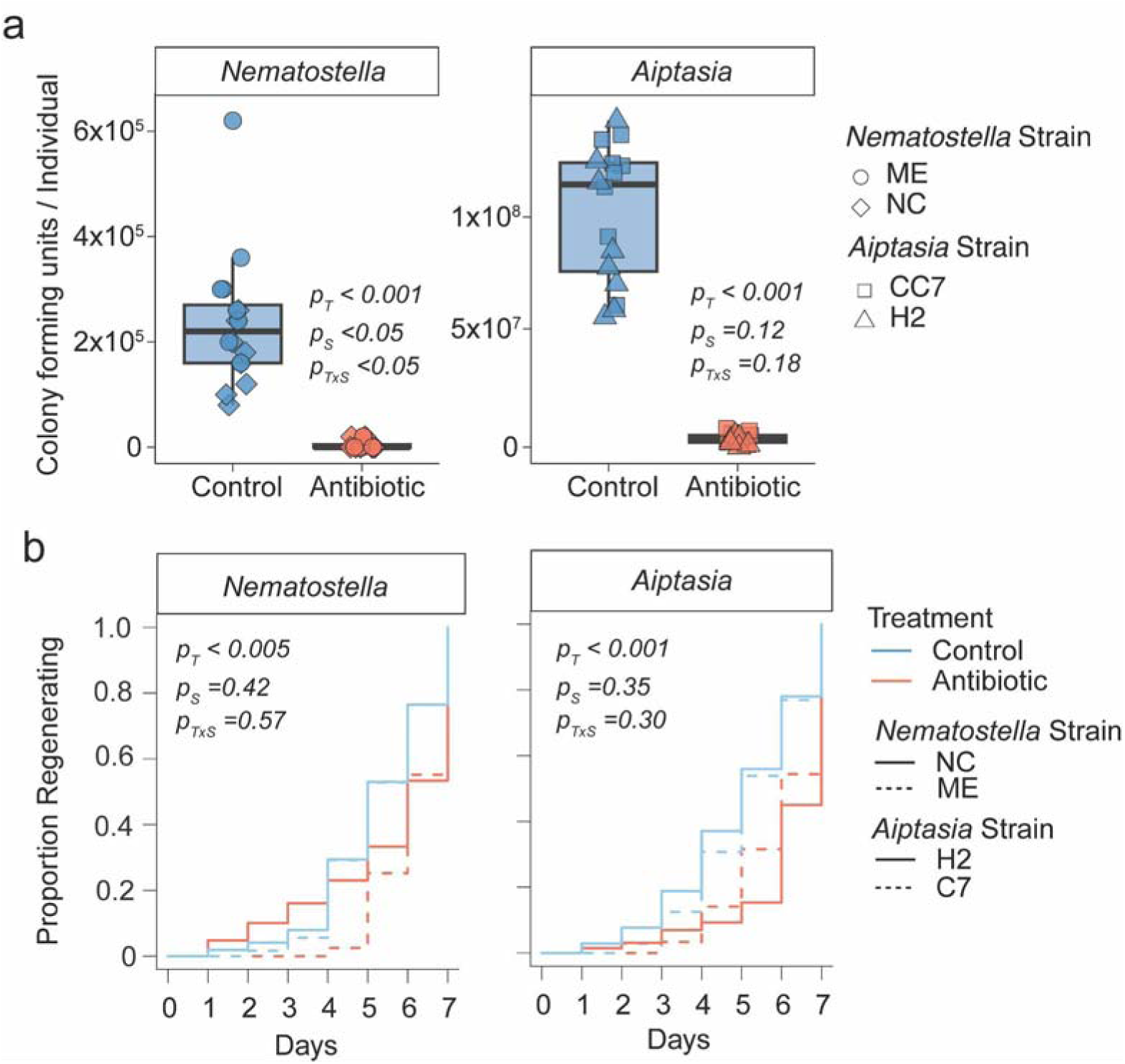
Anemone responses to antibiotic treatment. (a) Colony forming units (CFU) per individual under control and antibiotic treatments. Antibiotic treatment significantly reduced CFU counts (p_treatment_<0.001). (b) Proportion of individuals regenerating bisected oral discs over time between treatments. Line types represent different strains: *Nematostella* strains ME (solid) and NC (dashed), and *Aiptasia* strains CC7 (solid) and H2 (dashed), while color represents control (blue) and antibiotic treatment (red).

### Delayed oral disc regeneration in antibiotic treatments

To assess the impact of antibiotics on regeneration, anemones were bisected and the aboral halves were monitored daily for seven days to track oral disc regrowth, which was determined via tentacle emergence. Under control conditions, *Nematostella* and *Aiptasia* regeneration was first observed after 3 and 4 days, respectively (Fig. 1b). However, antibiotic exposure delayed regeneration in both species. In *Nematostella*, the proportion of individuals regenerating at day 5 was lower in the antibiotic treatment than control, with a significant difference in overall regeneration rate (Fig. 2b, Wald test, p<0.001). Specifically, ME *Nematostella* showed greater regeneration delays under antibiotics, with only partial recovery by day 7, whereas NC *Nematostella* experienced less pronounced delays (Fig. S1). Similarly, antibiotic treatments in *Aiptasia* resulted in delayed regeneration (Fig. 2b, Wald test, p<0.001). The extent of delay also varied by strain: CC7 showed a moderate delay by day 4, while H2 exhibited a more pronounced initial delay that improved by day 7 (Fig. S1). Regardless of strain level differences in magnitude through time, regeneration delays under antibiotic treatment were observed across strains in both species (Fig. 2b).

### Antibiotic treatment altered asexual reproduction in *Aiptasia*

To assess antibiotic effects on *Aiptasia* asexual reproduction, we counted pedal lacerates (tissue that detaches from pedal disc and develops into new clone) after 7 days under control or antibiotic treatment. The number of clones produced via pedal laceration in the antibiotic treatment was significantly higher (1.625±0.23) compared to controls (0.4375±0.12) (Fig. 3, F(1,28)=19.00, p<0.001). No significant differences between strains or a treatment *x* strain interaction were observed.

**Fig. 3.**
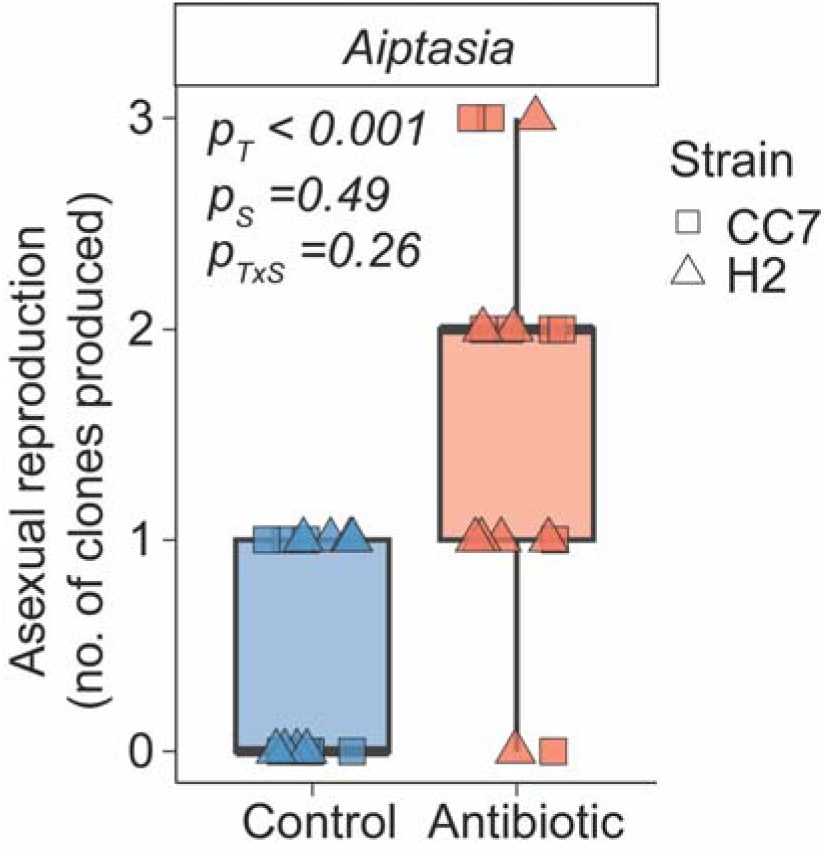
*Aiptasia*asexual reproduction, measured by numbers of clones produced via pedal laceration, between treatments. Shapes represent different strains: CC7 (square) and H2 (triangle), while color represents control (blue) and antibiotic (red) treatments. Note: no asexual reproduction was observed in *Nematostella*.

### Anemones developed fewer and shorter tentacles after exposure to antibiotics

To determine the effect of microbiome disruption on cnidarian tentacle phenotypes, we monitored tentacle number and length at the end of the experiment (day 7). Antibiotic treatment led to altered tentacle phenotypes (Fig. 4a). In *Nematostella*, tentacle numbers formed were reduced from 11.9±0.46 in control to 8.75±1.21 in antibiotic treatment (Fig. 4b, F(1,36)=6.57, p<0.05). *Aiptasia* exhibited a greater reduction in tentacle formation (52%) from 23.45±1.53 in control to 11.21±1.32 under antibiotic treatment (Fig. 4b, F(1,36)=40.62, p<0.001). No significant difference between strains was observed in either species (Fig. 4b). Similarly, tentacle length was significantly shorter in both species following antibiotic treatment. In *Nematostella*, tentacle length was reduced by approximately 60%, from 4.19±0.38mm to 1.56±0.19mm (Fig. 4c, F(1,47)=34.92, p<0.001) and *Aiptasia* showed an ∼80% reduction in tentacle length, from 4.99±0.39mm in controls to 0.97±0.10mm in antibiotic treatment (Fig. 4c, F(1,52)=156.47, p<0.001). Notably, H2 *Aiptasia* had longer tentacles overall than CC7 (Fig. 4c, p<0.001), which led to CC7 showing a less apparent reduction in tentacle length compared to H2 under antibiotic treatment (Fig. 4c, p<0.001).

**Fig. 4.**
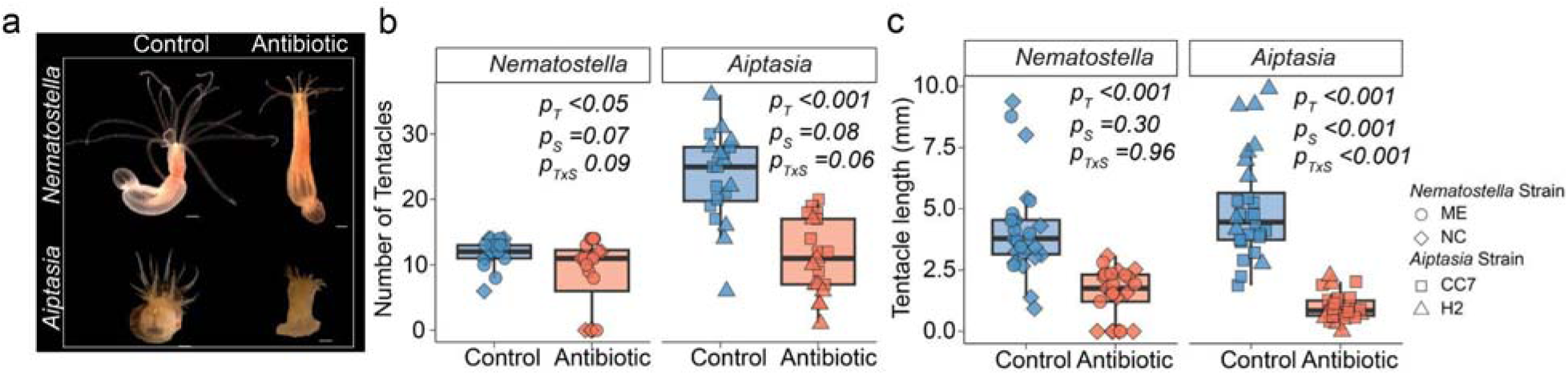
Phenotypic alteration in cnidarian tentacle phenotypes after exposure to antibiotic treatment. (a) Representative photographs of *Nematostella* (top) and *Aiptasia* (bottom) after 7 days of exposure to control (left) or antibiotic treatment (right). Antibiotic exposure resulted in shortened tentacles. (b) Average number of tentacles in different treatments (control=blue, antibiotic treatment=red). Antibiotic exposure resulted in lower tentacle regrowth relative to control. (c) Average tentacle length of individuals exposed to antibiotic treatment, which resulted in shorter tentacle lengths after 7 days relative to control anemones. Shapes indicate strains.

### Changes in the microbiome profiles of anemones exposed to antibiotics

To assess the impact of antibiotics on anemone microbiome profiles, we sequenced the 16S rRNA genes of whole anemones from one strain of *Aiptasia* (CC7) and *Nematostella* (ME). In *Nematostella*, antibiotic treatment resulted in a significant reduction of microbial alpha diversity compared to controls (Fig. 5a, *p<0.05*). Additionally, Principal Component Analysis (PCA) analysis revealed a distinct separation of bacterial communities between treatments (Fig. 5b, *p<0.001*), indicating that antibiotic exposure altered bacterial communities. Proteobacteria were the dominant group in *Nematostella*, and the composition within this phylum varied between treatments. Notably, the relative abundance of orders such as Burkholderiales and Francisellales decreased following antibiotic treatment, whereas Enterobacterales became more dominant (Fig. 5c), suggesting that antibiotic treatment differentially affected certain Proteobacteria taxa.

**Fig 5.**
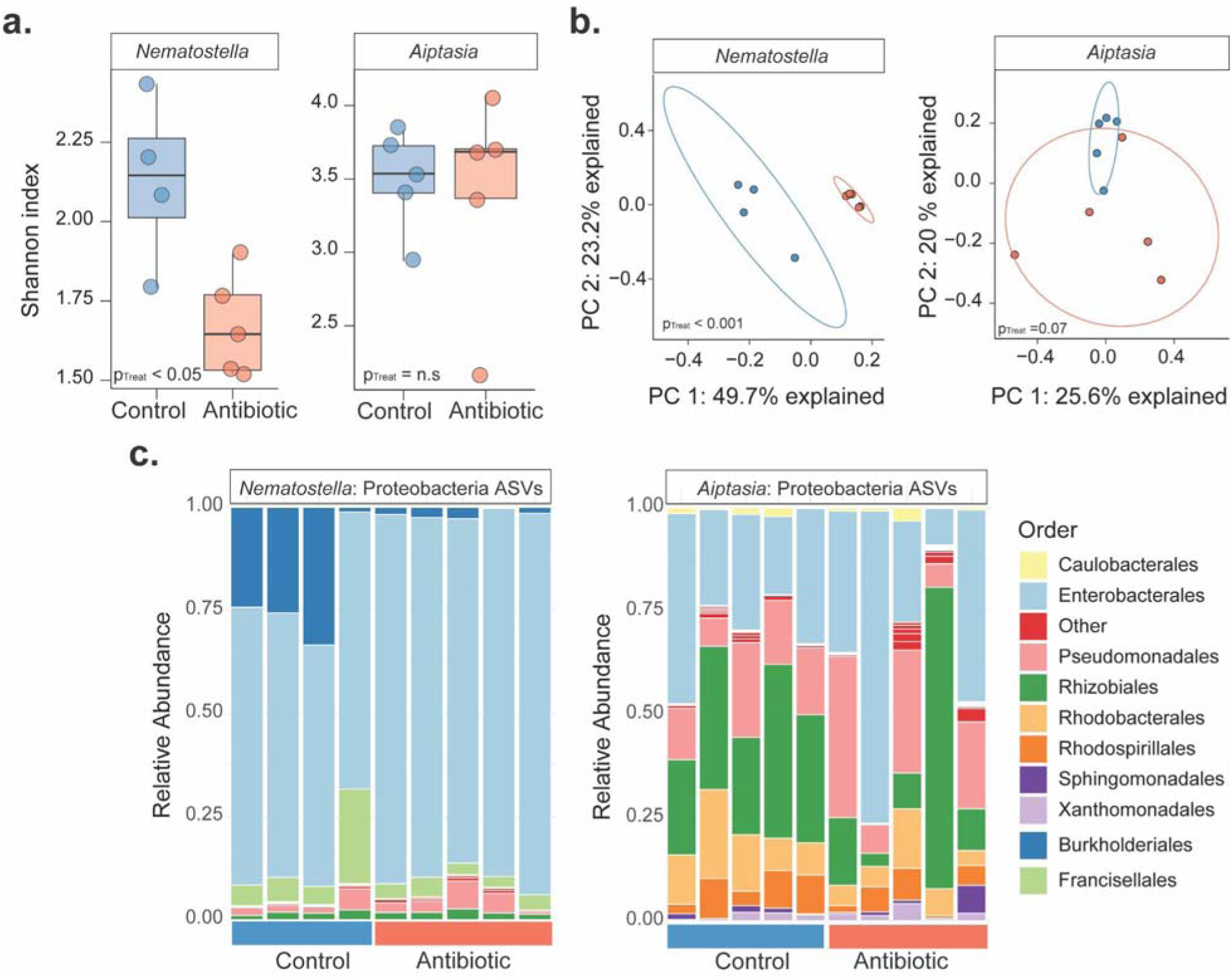
Microbial communities associated with *Nematostella* and *Aiptasia* exposed to antibiotic treatment. (a) Shannon diversity index comparing alpha diversity between treatments for *Nematostella* (left) and *Aiptasia* (right). A significant reduction in microbial diversity was observed in *Nematostella* (p<0.05) following antibiotic treatment, while *Aiptasia* showed no difference. (b) Principal Component Analysis (PCA) between treatments in *Nematostella* (left) and *Aiptasia* (right). *Nematostella* showed strong clustering between treatments (p<0.001), while *Aiptasia* exhibited muted clustering (p=0.07). (c) Stacked bar plots showing the relative abundance of Proteobacteria orders between treatments in *Nematostella* (left) and *Aiptasia* (right). Antibiotic treatment led to shifts in some members of Proteobacteria phylum in both cnidarians.

Conversely, antibiotic exposure in *Aiptasia* CC7 did not elicit a significant reduction in alpha diversity (Fig. 5a), which could be due to the inherently higher baseline microbial diversity of this species. PCA further showed that microbiomes of control and antibiotic-treated *Aiptasia* formed distinct clusters, although the separation was less pronounced than in *Nematostella*, with a marginally significant p-value (Fig. 5, *p=0.07*). This indicates that antibiotic treatment only had a moderate impact on overall microbial communities in *Aiptasia*. Exploring relative abundances of Proteobacteria in *Aiptasia* suggested that antibiotic treatment altered composition, although these changes were less pronounced than those observed in *Nematostella* (Fig. 5c). Specifically, within Proteobacteria, antibiotic treatment differentially influenced the microbial taxa in *Aiptasia*: the abundance of Rhodobacterales increased, whereas Xanthomonadales decreased.

## Discussion

We investigated the effects of antibiotic treatment on the regeneration capacity and associated microbial communities of two sea anemone models, *Nematostella* and *Aiptasia*. While both species experienced similar overall reductions in bacterial load, regeneration rate, and tentacle growth, their microbial community responses differed, suggesting that microbial stability and host-microbiome relationships vary across cnidarian species and may be linked to species-specific adaptations or algal symbiotic associations. In aposymbiotic *Nematostella*, overall phenotypic responses to antibiotic treatment were consistent between strains, suggesting the potential for a species-wide response to microbial disruption, although only two strains were explored. Conversely, the response of symbiotic *Aiptasi*a varied between strains, suggesting that genetic differences can influence resilience to microbial depletion. This variability highlights the need to account for genetic diversity in microbial resilience studies and suggests further research into the genetic factors influencing microbial community stability under stress.

### Antibiotic exposure impacts cnidarian regeneration and tentacle phenotypes

Here, antibiotic treatment reduced bacterial load in both cnidarian species, consistent with previous studies (Sweet et al. 2011; Bent et al. 2021; Costa et al. 2021; Connelly et al. 2023; Peng et al. 2023; MacVittie et al. 2024). Moreover, we found that antibiotic treatment and the resulting microbiome disruption delayed regeneration and had negative impacts on tentacle phenotypes in both cnidarian species, suggesting that host-associated microbiomes facilitate tissue regeneration and overall host physiology in these taxa. Strain-specific responses to antibiotics were not apparent in *Nematostella* (non-symbiotic) but were evident in *Aiptasia* (symbiotic). These differences could be attributed to baseline variation in the microbiomes of different *Aiptasia* strains (Herrera et al. 2017) or could reflect a more complex host-microbiome-symbiont (holobiont) interactions, where all components participate to maintain community structure and function under stress.

Notably, we observed an increase in pedal laceration in *Aiptasia* under antibiotic treatment, which contrasts (MacVittie et al. 2024) who found that exposure to antibiotics reduced pedal laceration. These contrasting results could be due to variation in the physiological state of the experimental *Aiptasia* or differences in experimental conditions, such as microbial community composition, antibiotic dose, or duration of exposure. However, pedal laceration is often a response to stress in *Aiptasia* (Hunter 1984; Clayton and Lasker 1985), which may be indicative of stress in regenerating animals exposed to antibiotics.

Regeneration requires significant energy and the presence of a beneficial microbiome (Nigro et al. 2014; Arnold et al. 2016). Therefore, disrupting beneficial microbes during the regeneration process could amplify organismal stress, leading to increased pedal laceration as *Aiptasia* are attempting to balance regeneration and microbial disruption. Microbial disruption following antibiotic treatment also aligns with observed delays in regeneration and tentacle phenotype reductions, which likely result from impaired microbial functions essential to various cnidarian biological processes (MacVittie et al. 2024), health (Ovchinnikova et al. 2006; Tinta et al. 2019), fitness (Lee et al. 2018; Weiland-Bräuer et al. 2020), and other developmental processes (Krueger et al. 2024). For example, antibiotic-induced microbiome depletion has been shown to delay the onset of strobilation (polyp-medusa transition) in jellyfish, impacting nematocyte development (Weiland-Bräuer et al. 2020; Peng et al. 2023). Antibiotic exposure has also been shown to delay *Nematostella* settlement time (Krueger et al. 2024). Alternatively, the impaired tentacle phenotypes that we observed under antibiotic treatment could be attributed, at least in part, to metabolic consequences caused by microbial depletion, such as mitochondrial dysfunction, nutritional deficiencies, and impaired immune functions, which disrupt cellular and physiological processes critical for development and growth (Moullan et al. 2015).

In addition to the observed effects on regeneration and tentacle phenotypes, microbial disruption may have broad consequences for immunity (Arnold et al. 2016) and metabolism (Ochsenkühn et al. 2018), both critical for maintaining homeostasis. Studies in other marine invertebrates suggest that the microbiome plays a key role in modulating immune system functions, with certain bacteria contributing to immune regulation and pathogen defense (Mannochio-Russo et al. 2023). Disruption of the microbiome could impair immune function, hindering wound healing and tissue regeneration (reviewed in (Eming et al. 2009; Karin and Clevers 2016)), and leaving the host more susceptible to pathogen infection. Furthermore, antibiotic treatments have been shown to alter metabolic profiles in mice, potentially disrupting energy allocation between growth, regeneration, and immune responses (VanHook 2018). Therefore, it is possible that the observed delayed regeneration and reduced tentacle number and length is the result of the combined effects of compromised immunity and metabolic processes due to microbial imbalances. While disentangling the direct effects of microbial associations on these host traits remains challenging, a deeper understanding of how each partner influences the symbiotic relationship is needed (Bove et al. 2022).

### Antibiotic exposure shifts microbiome compositions

16S microbial profiling revealed significant shifts in the diversity and composition of *Nematostella* microbiomes following the seven-day antibiotic treatment, highlighting the vulnerability of the microbiome to external perturbations. These disruptions likely contributed to the delayed regeneration seen in *Nematostella*, possibly due to the loss of beneficial microbial functions (Díaz-Díaz et al. 2021). In contrast, *Aiptasia* showed more stable microbial communities under the same antibiotic treatment, suggesting that their microbiomes are more buffered from these external stressors, perhaps through redundancy in the microbial community’s functions, where multiple microbes perform similar roles (reviewed in (McCauley et al. 2023)). Additionally, the symbiotic association of *Aiptasia* with Symbiodiniaceae, which maintain their own stable core microbiome (Lawson et al. 2018; Bernasconi et al. 2019), could provide a mechanism for microbiome resilience since algae are maintained inside host cells. This endosymbiosis may render removal of algal-associated microbes *via* antibiotic treatment more challenging in *Aiptasia*, and future work exploring the microbiome of both the host and algal symbiont in and out of symbiosis is warranted. Additionally, future work should investigate the effects of antibiotics on the photosynthetic apparatus of algal symbionts to disentangle the effects of symbiosis and to determine whether the ability of the algae to provide additional nutrition to the host is compromised under antibiotic treatment. These types of explorations would expand our understanding of the costs and benefits of symbiosis under antibiotic stress.

In *Nematostella*, antibiotic exposure resulted in reduced alpha diversity that was accompanied by clear reductions in microbial load. Such an effect can be caused by stressors that select for certain taxa while allowing more resilient ones to persist. This phenomenon could result in deterministic shifts in community composition (Zaneveld et al. 2017). Specifically, we observed strong microbiome shifts following antibiotic treatment, with notable decreases of Burkholderiales and Francisellales. These taxa are often associated with metabolic and physiological functions of the host. For example, Burkholderiales has been shown to support metabolic processes including nitrogen cycling (Baldani et al. 2000), which is important for cellular growth and tissue regeneration. Conversely, the shift in dominance of Enterobacterales in antibiotic-treated anemones may reflect its resilience, where it remains less affected than other taxa that fail to survive antibiotic treatment. Enterobacterales are not associated with beneficial roles in cnidarian development and are often known for their pathogenicity (Patterson et al. 2002; Krediet, Carpinone, et al. 2013). Their relative dominance following antibiotic treatment could lead to dysregulation of developmental processes, as these bacteria might not provide the same beneficial functions as suppressed bacterial groups. This shift could result in slower or impaired regeneration and developmental defects, as the microbiome’s overall functionality is altered.

While antibiotic treatment of *Aiptasia* caused less clear reductions in microbial diversity than *Nematostella*, microbial load was depleted and microbiomes still shifted. This reduction in microbial load, accompanied by a less pronounced reduction in diversity, suggests that although the overall microbial load was reduced in *Aiptasia*, the diversity of the remaining community was largely preserved. This pattern coupled with reduced regeneration under antibiotic treatment might imply impaired microbial function, where the taxa that survived antibiotic treatment are more likely to be resistant to antibiotics, but may lack the specific functions necessary for effective regeneration (Röthig et al. 2017; Cárdenas et al. 2022). Furthermore, the increased relative abundance of specific taxa like Rhodobacterales may be the opportunistic result of microbial members filling the functions of suppressed taxa. However, the persistent delay in regeneration suggests that these shifts do not fully compensate for the loss of essential microbiome functions. Additionally, the seven-day duration of antibiotic treatment may only partially capture the long-term effects on microbial communities and their implications for host physiology and regeneration. Lastly, 16S profiling was only conducted on strain CC7 of *Aiptasia* and it is possible that other strains exhibit divergent responses. Extending treatment times in future studies with additional Aiptasia strains could provide more comprehensive insights into the resilience of these communities and their critical roles in host health.

## Conclusion

We demonstrated that antibiotic treatment disrupts the microbiomes of two cnidarian models, which leads to impaired regeneration and reduction in overall health of *Nematostella* and *Aiptasia*. These shifts in microbial composition after antibiotic exposure underscore the susceptibility of the microbial communities of cnidarians to external factors, raising important questions about how cnidarians maintain resilience in the face of such disturbances. While *Nematostella* showed greater sensitivity to microbial disruption, *Aiptasia* exhibited strain-specific responses to antibiotic treatment, alongside an increase in pedal laceration—a stress response that further underscores the impact of microbial disruption. The more taxonomically stable, though depleted, *Aiptasia* microbiome suggests that host-microbe relationships vary between these cnidarians, and are likely influenced by the host’s evolutionary background or symbiotic associations (e.g., Symbiodiniaceae in *Aiptasia*). However, it is also possible that our seven-day treatment fails to fully capture the longer-term effects on microbial roles in host health. Future research on host-microbe interactions would benefit from examining how microbial shifts impact host system function (*i.e.,* metabolism, immunity) and how resilient these microbial communities are once the stressor is removed. Understanding these interactions may be useful for predicting how cnidarians will fare under changing environmental conditions, particularly in the context of climate change, where both microbiome stability and host health are at risk (Ziegler et al. 2019; Torda et al. 2020). As cnidarians continue to face environmental challenges, understanding their host-microbiome interactions will be important for developing strategies to ensure their survival.

## Acknowledgments

We thank Kian Thompson, Maria Valadez-Ingersol, and the BU Marine Program staff for experimental support; the BU Supercomputing Center for computational analysis assistance; Quinton Krueger and the Adam Reitzel Lab for the *Nematostella* strains; and Rachel Wright, the Philip Cleves Lab, and the Virginia Weis Lab for the Aiptasia strains.

## Supplementary Tables and Figures

**Fig S1.**
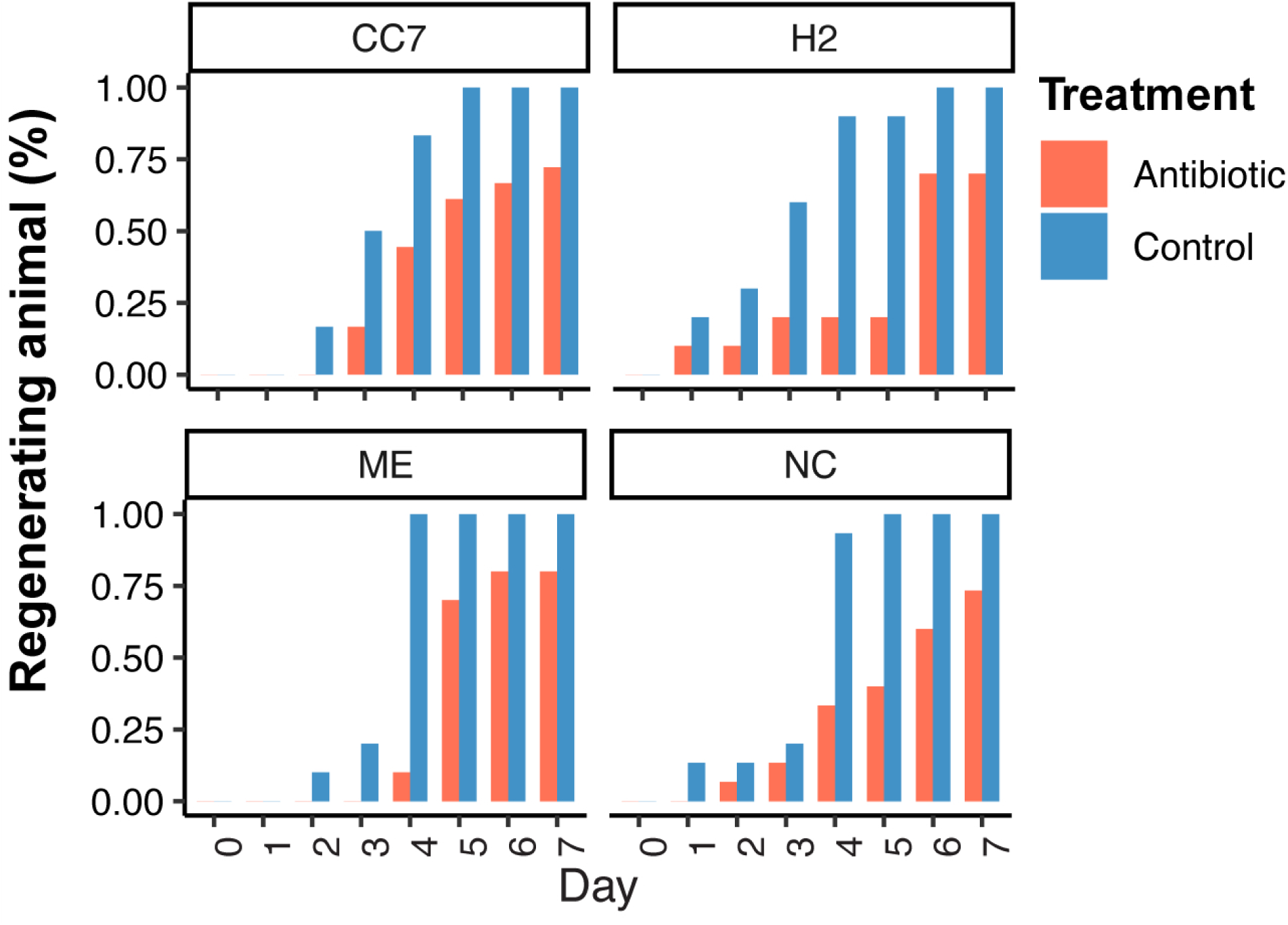
Percentage of regenerating animals over time in each strain: CC7 and H2 for *Aiptasia*, and ME and NC for *Nematostella*. Regeneration was tracked daily for 7 days, and the proportion of animals regenerating under antibiotic treatment (red) or control conditions (blue) is shown. Each treatment included 38 animals per strain and species.

